# Modelling the spread of resistance to *Varroa* in Australian feral honeybee populations

**DOI:** 10.1101/2022.02.10.479847

**Authors:** Robert Owen, Jean-Pierre Scheerlinck, Mark Stevenson

## Abstract

Australia is the only country where Western honeybees, *Apis mellifera*, are not infested with the mite *Varroa destructor*. Hence, a collapse in the feral honeybee population, brought about by an incursion and spread of *V. destructor*, would have serious consequences for Australian horticulture given its dependence on managed and feral honeybees for pollination. Managing *Varroa* in commercial colonies is well understood and can be achieved, although at significant bee-health costs, by using miticides. Protecting the feral population in the event of a *Varroa* incursion is much more difficult, but nevertheless imperative. One way to mitigate against collapse of the feral population is to seed it with *Varroa*-resistant queens, so as to accelerate the spread of resistance. We developed a simulation model of the spread of *Varroa*-resistance in feral honeybee populations following the introduction of *Varroa*-resistant queens into the managed population.

We show that, compared with a do-nothing scenario, seeding the managed honeybee population with *Varroa*-resistant queens was only effective in decreasing the length of time for the feral honeybee population to recover when the size of the managed resistant population was large compared to the size of the feral population. This situation may be achievable in some urban areas where numbers of managed colonies are high and habitat for feral colonies is limited.

## Introduction

Australia is the only country where the ectoparasitic mite *Varroa destructor* is not endemic in the honeybee population (Gross et al., 2020; Noel et al., 2020; Rosenkranz et al., 2010). Stringent border security and quarantine regulations are in place to minimize the risk of the mite entering Australia, although many believe an incursion is inevitable and that *Varroa* will eventually become endemic (Cook et al., 2007; Hafi, 2012). An incursion of *Varroa* is likely to have a serious impact on feral honeybee colony numbers. This, in turn, will have serious consequences for Australian horticulture given the dependence of these industries on both managed and feral bees for pollination. A 2003 study concluded that the value of bee-pollination to Australian horticulture was in the order of $2.86 billion USD (2019 USD equivalents) in 1999-2000 (Davis and Gordon, 2003). Another study concluded that thirty-five crops are either partly or completely dependent on bee pollination and that the value of horticulture in Australia in 2019 was $6.39 billion USD (2019 USD equivalents) (Senate, 2014). Globally, honeybees contribute around $265 billion USD (2019 USD equivalents) to the international economy (Gallai et al., 2009). This estimate does not take into account pollination of pastures such as clover or Lucerne used for cattle (Allsopp, de Lange et al. 2008). If the feral bee population collapsed in Australia there would be an immediate fall in the value of horticulture exports as well as the country’s ability to provide food to feed its local population. To illustrate the importance of feral honeybee colonies to horticulture with an example, the production of macadamia nuts require seven colonies per hectare for optimum pollination (Keogh, 2010) but commercial beekeepers only supply 2.5 hives per hectare (private correspondence, Macadamias Australia). The substantial shortfall of 4.5 colonies per hectare is made up by feral colonies.

Controlling *Varroa* infestations in managed colonies is well understood (Caron, 2015; Delaplane et al., 2005; Roth et al., 2020). Although complicated by the price of Mānuka honey, when *Varroa* was detected in New Zealand in 2000, the price of both honey and pollination hives increased which offset the added costs of managing the mite (Sommerville, 2008). Since much of Australian horticulture is dependent on feral colonies for pollination it is imperative that authorities take steps in advance of a *Varroa* incursion to protect the feral honeybee population (Hafi, 2012). To offset this risk biosecurity authorities in Australia should: (1) continue with current border security arrangements (Australia, 2016; BeeAware, 2019); (2) acknowledge that an incursion is highly likely and develop detailed, fit-for-purpose post incursion consequence mitigation measures; and (3) develop pre-incursion strategies to reduce the severity of *Varroa* spread, if and when an incursion occurs.

A widely used control measure for *Varroa* is to introduce miticides into the hive at concentrations that kill mites but not bees. Miticides, however, have sub-lethal effects that harm bees (Mullin et al., 2010; Noel et al., 2020; Roth et al., 2020). A conceptually attractive alternative method is to breed bees that are resistant to the mite (Büchler et al., 2010; Rinderer et al., 2010). Two strains of bees that show high resistance are the *Varroa* Sensitive Hygienic line, VSH (Danka et al., 2015) and Russian bees (Rinderer et al., 2010; Westra, 2007). Both strains are commercially available in the USA and Europe but not in Australia as importation of bees is subject to strict quarantine protocols. In Australia, there is a need to understand the effect of introducing resistant queens into the managed population with the aim of increasing resistance in the feral population by swarming and resistant drones mating with feral virgin queens. In this context, important considerations are: (1) can resistance genetics be introduced into the feral population; (2) can high levels of *Varroa*-resistance be introduced into the managed and feral populations prior to *Varroa* becoming endemic; and (3) can a scenario be formulated that will speed-up the increase in feral resistance post-*Varroa* incursion (van Alphen and Fernhout, 2020)?

To address these issues, we developed a simulation model to estimate the likely effect of using managed colonies to seed the feral population with *Varroa*-resistant alleles as a way of accelerating the growth of resistance in the feral population.

## Materials and methods

### Honeybee reproduction

Honeybee reproduction is unusual in that a virgin queen, a few days after hatching, goes on one to three mating flights and copulates with 16 to 20 random drones from other colonies. During the mating process it is unlikely that the virgin queen will mate with drones from her own colony since the virgin queen flies a longer distance from her home colony to a drone congregation area (DCA) compared with drones (Koeniger et al., 2014; Seeley, 1978; Seeley, 2010; Seeley, 2019; Winston, 1991, 2014). The queen will store one to four million sperm in her spermatheca (Koeniger and Koeniger, 2008) and will not mate again during her two to four year lifetime. At peak colony-growth, usually in the spring and early summer, the queen lays around 1,500 eggs per day to maintain the colony population at around 20,000 to 40,000 female worker bees. During development in the queen’s ovaries a single set of chromosomes will be included in each egg. As the egg passes down the oviduct to the vagina it may be fertilized by sperm from one of the 16 to 20 drones which mated with the queen resulting in the egg becoming diploid and developing into a female worker. Non-fertilized eggs develop into haploid male drones.

Reproduction at the colony level occurs as a result of swarming. Each year approximately 80% of feral colonies (Winston, 1991) and 10% to 30% of managed colonies swarm (S Williamson, personal communication). Swarming is a process by which approximately half the bees, together with the old queen, leave to establish a new colony. Around fourteen days before a swarm event occurs, the old queen lays about six to eight female eggs that are destined to become new queens; these hatch two or three days after the swarm leaves the colony. The first virgin queen to hatch usually kills the other, unhatched, queens. Since the queen only goes on mating flights to DCAs during the first few days of her adult life, there is only a limited amount of sperm in her spermatheca. This becomes depleted in two to four years at which time the female workers in the colony will replace her with a new queen in a process called supersedure (Crane, 2015). supersedure is similar to swarming in that a set of new virgin queen pupae are reared, one of whom will survive to head the colony. With supersedure the colony does not swarm but remains in the old nest. The old queen is believed to be removed by workers later (Koeniger et al., 2014; Seeley, 1978; Seeley, 2010; Seeley, 2019; Winston, 1991, 2014).

### Construction of model

The model was designed to identify the correlation between feral population size for 15 years after *Varroa* becomes endemic and key input parameters, including: (1) the proportion of resistant queens introduced into the managed population each year; (2) the relative size of the managed honeybee population to the feral population; (3) increasing the number of drones produced in resistant managed colonies; (4) the proportion of resistant drones and workers in a colony; and (5) the probability of colony death conferred by the possession of *Varroa* resistant genetics. The two parameters that can be altered by beekeeper action are the proportion of *Varroa* resistant queens in the managed population and the number of drones produced by each resistant managed colony.

The model of the feral population is more complex than for the managed population because it includes colony death as a result of *Varroa* and/or lack of resources such as food, water, or nesting sites. Swarms form the managed population also join the feral population. In the model, the feral and managed populations were modeled as two semi-independent sub-populations that have different characteristics and genetic structures, shown in Figure 1.

**Figure 1.**
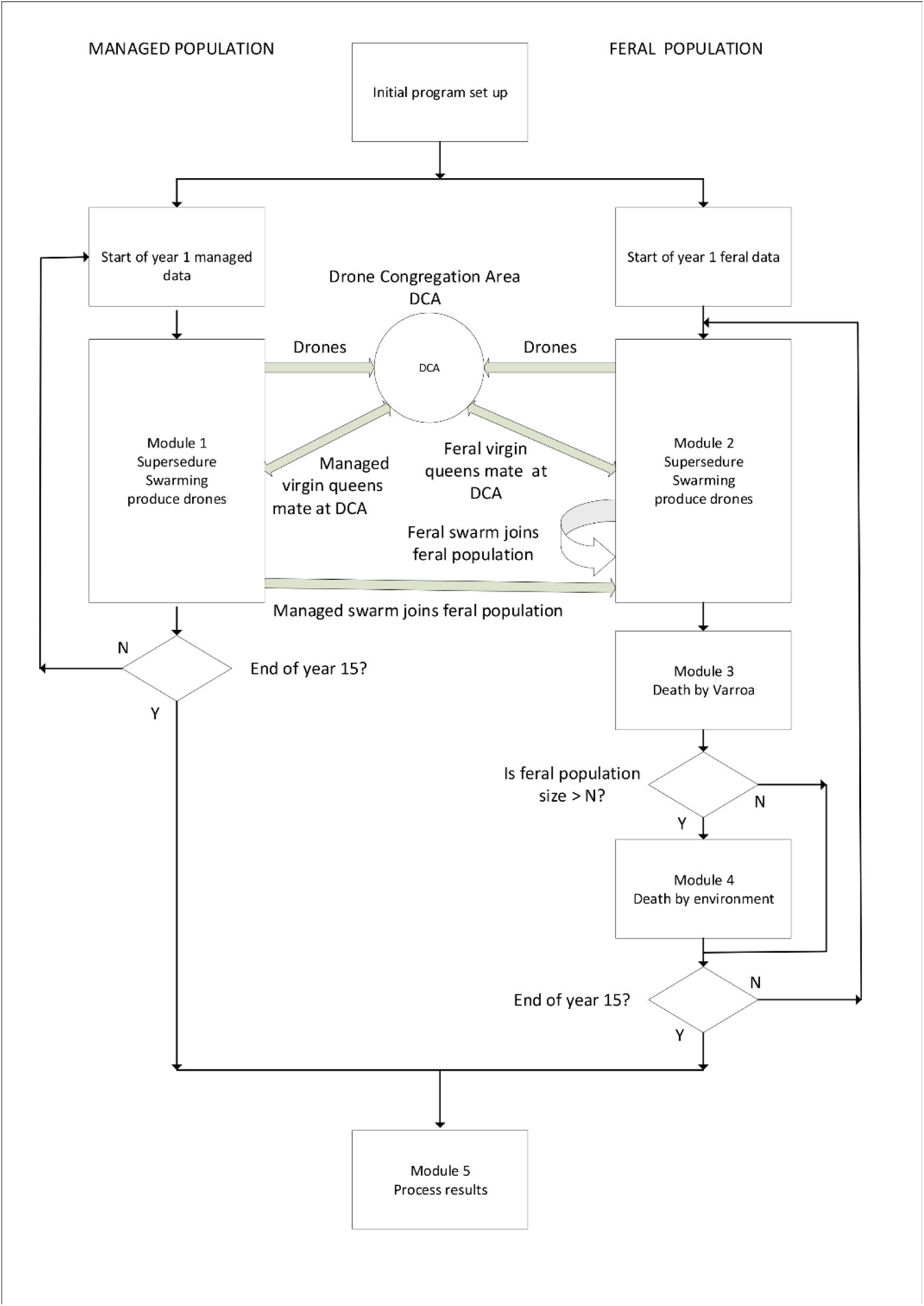
High-level diagram showing the relationship between managed and feral populations. See Supplementary Material 1 for detailed description.

#### Assumptions

As a simplification our model assumes that managed and feral colonies are evenly mixed even though commercially managed colonies, in Australia, are more likely to be managed in clusters (called apiaries) and moved about every three weeks during the pollination and honey producing season. A second assumption is that *Varroa* is endemic in the honeybee population and evenly distributed among managed and feral colonies. This is a simplification since *Varroa* will spread into a population over a number of years.

A queen is assumed to possess either none, one, or two *Varroa* resistance alleles on her two chromosomes. The queen’s spermatheca contains sperm collected during her mating flights, a proportion of which may also contain *Varroa*-resistance alleles. The female worker larvae may therefore possess none, one or two resistance alleles depending on whether the egg contained a resistance allele from the queen and whether the sperm used to fertilize that egg carried a *Varroa*-resistant allele. Our model assumes that resistance is carried by a single or a set of closely-spaced alleles that remain together during meiosis. It is likely however that resistance arises from the presence of several alleles, sometimes coding for independent traits (Boecking et al., 2000; Lapidge et al., 2002; van Alphen and Fernhout, 2020). *Varroa*-resistance can take many forms, for example the Varroa Hygienic Sensitivity trait, VHS, is recessive (van Alphen and Fernhout, 2020) and additive (Pers. comm. Bob Danka). While our model assumes that resistance alleles are additive, sensitivity analysis shows that the conclusions would be the same if the *Varroa*-resistance alleles are dominant, co-dominant, or recessive

#### Construction of Model

See Supplementary Material 1 for detailed description of model. As shown in Figure 1, the model is comprised of three ‘compartments’: (1) the feral population; (2) drone congregation areas (comprised of a mixture of managed and feral honeybees); and (3) the managed population. Within each compartment we estimate colony population turnover, keeping tally of *Varroa*-resistance and its consequent protective effect on colony mortality from *Varroa*. At the start of each 15-year simulation the proportion of managed colonies that contained a *Varroa*-resistance allele was varied to simulate the effect varying levels of requeening with resistant queens. This proportion was reset to the start of year-1 managed population levels at the beginning of each year during a 15-year simulation. Requeening every year is practiced by commercial beekeepers using purchased queens since queens less than a year old lay more eggs and are less likely to swarm; re-setting the proportion of colonies that contained a *Varroa*-resistance allele accounted for this practice. The number of *Varroa*-resistant feral colonies was set to 5% at the start of year 1 since studies in the USA and Europe have shown that if a feral honeybee population is not to be eliminated completely as a result of *Varroa*, around 5% of feral colonies need to survive the initial infestation (Fries and Rosenkranz, 2006; Kefuss, 2016; Locke and Fries, 2011; Seeley et al., 2015; van Alphen and Fernhout, 2020).

Many of the processes used in the three modules are stochastic. These include the number of *Varroa*-resistant drones a virgin queen mates with, how many resistance alleles a new queen receives, which colonies are killed by *Varroa* and/or lack of resources, as well as the number of managed *Varroa*-resistant colonies. Details of each of the stochastic variables included in the model including their distribution, input parameters and references to justify the values used are provided in Table 1.

**Table 1.** Stochastic parameters included in the model, their distributional form and supporting references.

| Parameter | Distribution | References |
| --- | --- | --- |
| Number of feral colonies in sub-population. | Uniform (500, 3000). This corresponded to a fixed managed population of 1,000 colonies with the feral population being 0.5 to 3.0 the size of the managed population. | (Oldroyd, Thexton et al. 1997, Arundel, Oxley et al. 2014, Hinson, Duncan et al. 2015) |
| Proportion of managed colonies with resistant queens | Uniform (0, 1.0) | Commercial beekeepers requeen most of their colonies every one or two years. |
| Resistance bands for each colony, $u, v, w$ | $u, v \wedge w$ are randomly selected at the start of each 15-year simulation such that:<br>$0 < u < v < w < 2$ | The average number of resistant alleles possessed by workers in a colony can vary between 0 and 2. No resistance in the colony to each worker possessing an allele on each chromosome. |
| Probability of death for a colony, $x, y, z$ | $x, y \wedge z$ are randomly selected at the start of each 15-year simulation such that:<br>$0.05 < x < y < z < 0.95$ | A colony in the low resistance band $u$ would have a high probability of death $z$. $y$ pairs with $v$ and $x$ pairs with $w$. |
| Colonies selected for "death" by *Varroa* | At the end of each year, a proportion $x$ of colonies in the low resistance band $u$ would be removed from the model. Colonies are randomly selected for removal. Same for $y, v$ and $z, w$. | |
| Colonies selected for death due to carrying capacity of environment | When the feral population exceeds the carrying capacity of the environment, $N_f$, excess colonies are removed from the model until the population reaches $N_f$. Colonies are randomly selected for removal at the rate $(x, u), (y, v)$ and $z, w$. | It is assumed that colonies that have less resistance to *Varroa* will be more likely to die from lack of resources than more resistant colonies. |
| Drones produced by resistant managed colonies | Uniform (0.5, 3.0). Resistant colonies produce between 0.5 and 3.0 the number of drones produced by non-resistant colonies. | Managed colonies can be encouraged to produce more drones by placing drone comb in the brood box. |
| Percentage of resistant queens in feral population | Uniform(0,0.1) |  |

When the number of alleles conferring *Varroa*-resistance carried by a queen (0, 1 or 2) is known and the proportion of sperm in her spermatheca that carry the allele has been estimated, the proportion of resistant workers in her colony can also be calculated. Depending on the proportion of worker bees that possess resistance, the colony is stochastically assigned a probability of death for that year. The larger the proportion of resistant workers the less likely the colony will die, death usually occurring during late autumn or winter (Rosenkranz et al., 2010; Roth et al., 2020). Colonies with a higher number of resistant workers are also more likely to spread resistance genetics as a result of swarming or by producing drones that mate with virgin queens in DCAs (Neumann and Blacquiere, 2017).

#### The managed population

The left-hand side of Figure S1 shows steps in the simulation of the managed population over a one-year season, and is repeated 15 times to simulate 15-years, module 1. It is assumed that managed colonies that die are replaced with new colonies by the beekeeper. A portion of the managed population is requeened every year with *Varroa*-resistant queens purchased or reared by beekeepers. It is also assumed that all swarms from managed colonies join the feral population, while virgin queens’ mate and return to their original hive. The managed population adds resistance to the feral population both by the production of *Varroa*-resistant drones that fly to DCAs to mate, as well as *Varroa*-resistant queens that join the feral population as a result of swarming.

The genetic structure of the managed population changes over the year influencing the genetic make-up of drones in the DCAs. However, this change is not used as an input for the following year since most managed colonies are requeened each year (as described above) resetting the genetic structure of the managed population to the start of year-1 values. Thus, apart from some random changes during the year, the structure of the managed population remains the same each year of a 15-year simulation.

#### The feral population

The right-hand side of Figure S1 shows steps in the simulation of the feral population during a year, modules 2 to 4. At the start of year 1 the initial structure of the feral population is entered into the model from which a 10-field data structure (Table S1, Supplementary Material 1) for each colony is constructed. Both feral colonies and managed colonies produce drones that fly to DCAs to mate with feral and managed virgin queens. These drones may carry with them resistance to *Varroa* which they pass to the virgin queen at the time of mating. Some colonies also supersede and replace their queen; some colonies will also swarm, producing a virgin queen that flies to the DCA to mate before returning to the same colony. When a colony swarms, approximately half the bees leave to establish a new feral colony, leaving the new virgin queen to head the old colony in the same nest. Since selection kills fewer resistant colonies the proportion of drones carrying *Varroa*-resistance increases over the 15-year simulation period.

#### Feral population deaths by Varroa and lack of resources

Figure 1 provides further detail of the simulation of the feral population over late autumn and winter. In module 3, *Varroa*-resistance of each colony is calculated, and a probability of death is stochastically assigned. Depending on the calculated probability of death, the model randomly selects colonies that will not survive to the following year due to death by *Varroa*.

When colony death by *Varroa* has been carried out, the number of surviving feral colonies is compared with the colony carrying capacity of the environment, *N*_*f*_, module 4. If the number of feral colonies is greater than *N*_*f*_ the weakest colonies are removed, reducing the feral population to the size that it was at the start of year 1. It is assumed that the colonies to be removed for carrying-capacity reasons are those that have been weakened by *Varroa*. Our reasoning here was that if a colony was already weakened by *Varroa* it is more likely to die if resources are scarce. Feral colony carrying capacity is assumed to be equivalent to the number of pre-*Varroa* feral colonies. We assumed that if the environment could support more than the pre-*Varroa* number, there would have been less environmental causes of colony loss prior to *Varroa* and the feral population would have expanded to the maximum number able to be supported Seeley, 2006, 2017; Seeley et al., 2015; Utaipanon et al., 2020).

The output from module 4, the end of year structure of the feral population, is passed back to the start of module 1 and used as input for the following year. When the model has estimated the growth of resistance over 15 years, the simulation is ended, module 5.

See Supplementary material 1 for a detailed explanation of the model.

#### Calculation of resistance to *Varroa*

Each feral colony is assigned to one of three *Varroa* resistance bands, low (*x*), medium (*y*) or high (*z*), depending on the number of resistant alleles within the colony, such that 0< *x* < *y* < *z* <2. Codominance results in the upper bound being 2.0 because each chromosome can carry a resistance allele. Each of the three resistance bands was assigned a yearly probability of colony death *u, v* or *w*, respectively with 0.05 < *u* < *v* < *w* < 0.95. Colonies in the low resistance band *x* have a higher probability of death *w* than those in a high resistance band, while the probability of swarming (0.3) was lower in the low-resistance band compared to the medium (0.5) and high-resistance bands (0.8). At the start of each 15-year simulation, both the width of the resistance bands and the probability of death were randomly varied.

At the start of each 15-year simulation the size of the feral population was assigned by taking a random draw from a uniform distribution with a minimum value of 500 and a maximum value of 3000 (Table 1) with this range estimated to be between half and three times the size of the managed population in the same area (1,000) (Arundel et al., 2014; Hinson et al., 2015; Oldroyd et al., 1997). The number of managed colonies in eastern Australia during 2018 and 2019 was estimated to be 596,000 (BeeAware, 2019), although only a subset of these were modelled.

Two scenarios were simulated, as described in Table 2. For the first it was assumed that a random number of managed colonies would be seeded with resistant queens in year 1 and this addition of resistant queens would be maintained each year by beekeepers for the 15-year simulation period. The second scenario was identical to the first with the additional intervention where beekeepers would be encouraged to produce more drones by placing drone-embossed comb and resistant queens into their hives to increase the percentage of *Varroa*-resistant drones in the mating population.

**Table 2.** Details of the *Varroa* management scenarios presented in this paper that were investigated as possible way to increase resistance in the feral population.

| Scenario | Details |
| --- | --- |
| 1 | Proportion of managed colonies seeded with *Varroa* resistant queens. Requeening colonies with resistant queens is the most likely method of introducing resistance into the managed population. The proportion of managed colonies seeded varied between 0% and 100%. |
| 2 | Number of drones produced by each managed colony headed by a *Varroa* resistant queen. The number of drones produced by each resistant managed colony varied between 0.75 and 3.0 the number produced by non-resistant colonies. Drone numbers can be increased by adding drone comb to hives. |

For each of the two scenarios the model was run multiple times, for 20,000 15-year simulation periods. In total, 500,000 15-year simulations were run. Model outputs are presented using descriptive statistics for the proportion of the original population of managed and feral colonies present at the end of the 15-year simulation period and the smallest proportion of the original feral colony population present before recovery. The number of feral colonies and the percentage of resistance queens introduced into the population at the start of the simulation period, the percentage of resistant queens in the feral population at the start of the simulation period and the number of feral colonies relative to managed colonies were correlated (Hauke and Kossowski, 2011; Rodgers and Nicewander, 1988; Xu et al., 2016).

The model is described in detail in Supplementary Material 1, while the Python code is shown in Supplementary material 2.

The model was constructed using Python version 3.8.5 (Chudoba et al., 2013; Python, 2019f) using the packages Numpy (van der Walt et al., 2011), random (Matsumoto and Nishimura, 1998; Python, 2019b), copy (Python, 2019c), csv (Python, 2019d), time (Python, 2019e), matplotlib (Matplot, 2019), and math (Python, 2019a). See Supplementary Material 2 for Python code.

## Results

Table 3 provides descriptive statistics of the proportion of the original feral colony population present at the end of each of the 20,000 15-year simulations and the smallest proportion of the original feral colony population present before recovery. For most combinations of variables, the feral population collapses and does not recover within 15 years. In some scenarios, however, where the death rate (*z*) is high if there is little resistance (band *u*), and lower (*x, y*) in medium to high resistance (bands *v* and *w*), the feral population recovers after 10 to 12 years.

**Table 3.** Descriptive statistics of: (a) the proportion of the original feral colony population present at the end of each of the 20,000 15-year simulations; and (b) the smallest proportion of the original feral colony population present before recovery.

| Scenario | Outcome | Initial population (*n*) | Proportion of original population remaining |  |  |
| --- | --- | --- | --- | --- | --- |
|  |  |  | Mean (SD) | Median (Q1, Q3) | Min, Max |
| 1 | $N_m$ at end of 15 yrs | 1,000 | | | |
| | $N_f$ at end of 15 yrs | Uniform (250, 3,000) | 0.448 (0.321) | 0.324 (0.191, 0.673) | 0.054, 1.0 |
| 1 | $N_m$ at minimum | 1,000 | | | |
| | $N_f$ at minimum | Uniform (250, 3,000) | 0.281 (0.158) | 0.24 (0.167, 0.357) | 0.054, 0.902 |
| 2 | $N_m$ at end of 15 yrs | 1,000 | | | |
| | $N_f$ at end of 15 yrs | Uniform (250, 3,000) | 0.4478 (0.3213) | 0.324 (0.191, 0.673) | 0.054, 1.0 |
| 2 | $N_m$ at minimum | 1,000 | | | |
| | $N_f$ at minimum | Uniform (250, 3,000) | 0.281 (0.158) | 0.24 (0.167, 0.357) | 0.054, 0.902 |

SD: standard deviation; Q1: first quantile; Q3: third quantile.

Table 4 lists the correlation between input parameters and the size of the feral population at end of year15. Even though there was a small correlation between the percentage of *Varroa*-resistant queens introduced into the managed population at simulation start and final feral population size, there was a larger correlation between the size of the managed population compared to the feral population. The smaller the size of the managed population relative to the feral population, the lesser the effect managed *Varroa*-resistant queens had on the feral population. Resistance within the feral population at the start of a simulation had the largest effect on the final feral population size. The probability of death for the medium *Varroa*-resistance band (parameters *y* and *v* in the model) had the strongest, negative, association with final feral population size.

**Table 4.** Pearson’s correlation coefficients (and their 95% confidence intervals) quantifying the correlation between selected input parameters and the size of feral population at the end of a 15-year simulation period (scenario 1).

| Parameter | Pearson’s rho (95% CI) |
| --- | --- |
| Percentage of resistant queens in managed population at start | 0.033 (0.020 to 0.046) <sup>a</sup> |
| Percentage of resistant queens in feral population at start | 0.288 (0.276 to 0.300) |
| Size of feral population compared with managed population | 0.295 (0.283 to 0.307) |
| Probability of death, low resistance | -0.416 (0.405 to 0.427) |
| Probability of death, medium resistance | -0.656 (0.649 to 0.663) |
| Probability of death, high resistance | -0.383 (0.372 to 0.394) |

<sup>a</sup> Interpretation: The correlation between the proportion of resistant queens in the feral colony population at the start of the simulation and the size of the feral population after a 15-year simulation period was 0.033 (95% CI: 0.020 to 0.046).

Table 5 lists the correlation between the use of drone comb in managed hives and final feral population size. Use of drone comb in managed colonies had little effect on minimizing the collapse of the feral population. Drone comb had a small positive effect on the recovery of the feral population after 15 years.

**Table 5.** Pearson’s correlation coefficients (and their 95% confidence intervals) quantifying the correlation between increased drone production in resistant, managed colonies (scenario 2) and the minimum size of the feral colony population after collapse and the size of the feral population at the end of a 15-year simulation period.

| Parameter | Pearson’s rho (95% CI) |
| --- | --- |
| Minimum size of the feral colony population after collapse | 0.115 (0.102 to 0.128) <sup>a</sup> |
| Size of the feral colony population at the end of year 15 | 0.168 (0.155 to 0.188) |
| Year of first increase in population size after collapse vs. % managed resistant queens | -0.113(-0.126, -0.1) |

<sup>a</sup> Interpretation: The correlation between the increase in drones from resistant colonies in the managed population to the minimum size of the feral population after its collapse was 0.037 (95% CI: 0.023 to 0.050).

Our results show that the number of *Varroa*-resistant queens introduced into the managed population had little effect both on the minimum size of the feral population after its collapse, as well as its size at the end of 15 years. The only exception to this was if the resistant managed population was larger than the feral population, which, in most situations, will be an unlikely scenario (Arundel et al., 2014; Hinson et al., 2015; Oldroyd et al., 1997).

The managed population produced drones and swarms that have a genetic structure that was not changed over the 15-year simulation period. During a simulation, usually up to about year 5, when the managed population contained a higher proportion of *Varroa*-resistance alleles than the feral population, *Varroa*-resistance alleles were passed to the feral population. As selection progressed, the feral population contained a higher proportion of resistant alleles than the managed population. When the proportion of resistant alleles in the feral population was greater than that in the managed population, the managed population reduced the effects of feral selection by passing more non-*Varroa*-resistant alleles to the feral population. The larger the size of the managed population relative to the feral population, the greater this effect.

Figures 4a and 4b show the number feral colonies surviving, after death by *Varroa* and lack of resources or nesting sites as a function of simulation year. Shown is the collapse of the feral population by the end of year 4, followed by complete recovery by year 12. This result, experienced in a small number of *Varroa* infested locations globally, occurs when there is a high probability of death for low *Varroa*-resistant colonies and a low probability of deaths for medium and high resistant colonies (De Jong and Soares, 1997; Fries and Rosenkranz, 2006; Kefuss et al., 2004; Le Conte et al., 2007; Locke, 2016; Locke and Fries, 2011; Moretto et al., 1995; Seeley, 2006; Seeley et al., 2015; van Alphen and Fernhout, 2020). Figure 2b shows the relative proportion of feral colonies with high resistance, medium resistance and low resistance as a function of simulation year. In regions where the honey bee population has recovered within a few years (Africa and South America), about 30% of the feral population possessed resistance before the mite became endemic, while only about 5% to 7% of the population possessed resistance in areas where the feral population collapsed and did not recover(van Alphen and Fernhout, 2020).

**Figure 2:**
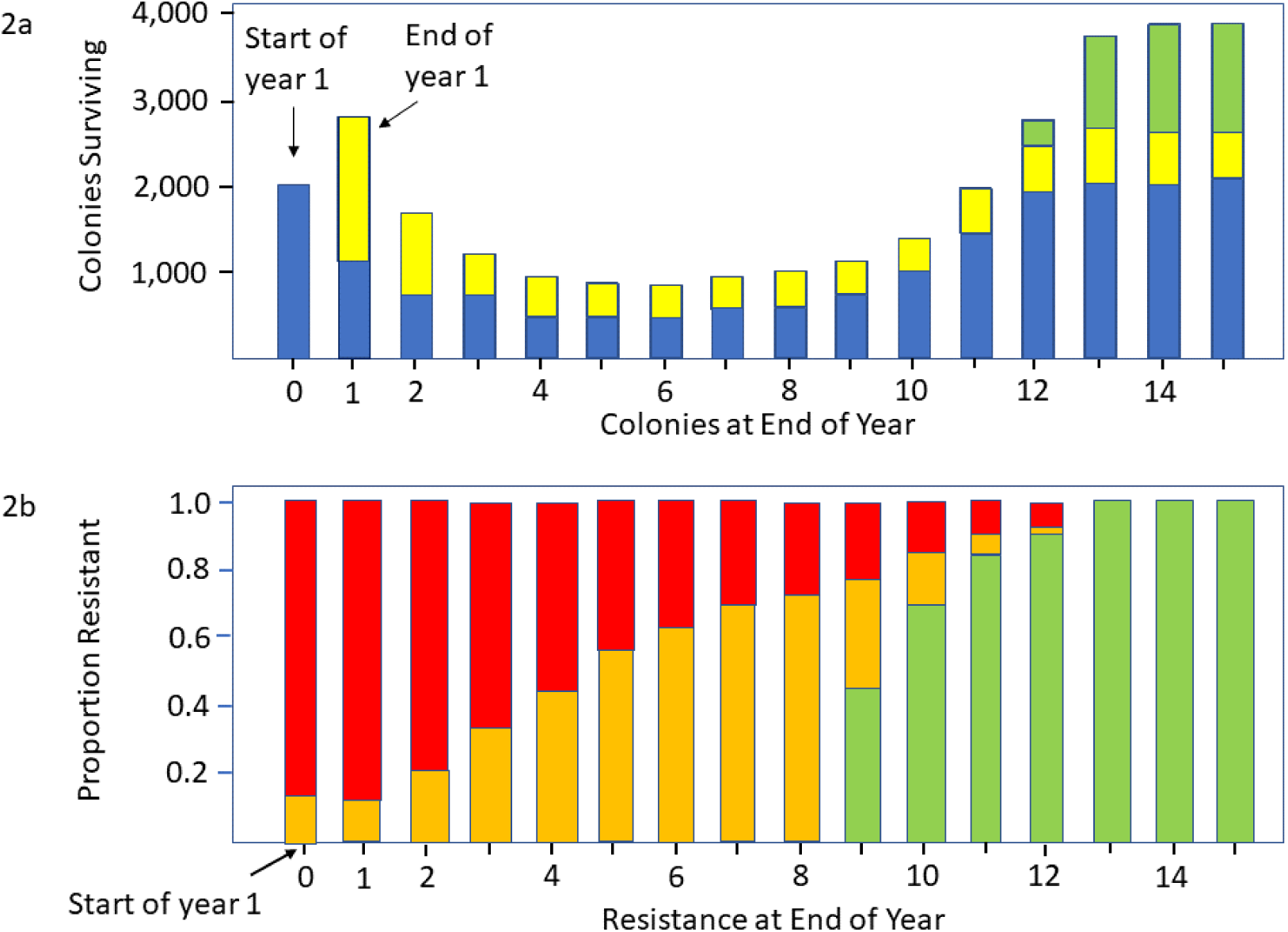
Stacked bar charts showing: Fig. (a) the number of colonies surviving 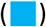, killed by *Varroa* 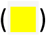 and killed by the environment 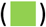 as a function of simulation year. Fig. (b) the percentage of feral colonies with high 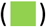, medium 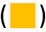 and low 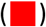 resistance as a function of simulation year. At start of year 1, 5% of feral colonies display medium resistance, 95% display low resistance. In this scenario, the probability of death by *Varroa* is low for colonies with medium levels of resistance, enabling the population to return to pre-*Varroa* numbers by the end of year 12. See text for details.

Parameter values that were used for the model results presented in Figure 2 are: number of managed colonies: 1,000; feral colonies: 2,000; resistance in feral and managed population at start of year 1: 5%; resistance bands [*x* = 0 to 0.4, *y* = 0.4 to 1.2, *z* = 1.2 to 2.0], probability of death [*u* = 0.9, *v* = 0.2, *w* = 0.1].

Figures 5a and 5b show complete collapse of the feral population with no signs of recovery after 15 years. One set of parameters that replicates this is given below, although other combinations lead to the same effect. Generally, if resistance confers little to a reduced probability of death the population will collapse. This is the situation in most infested regions globally, with only a few isolated colonies surviving.

Parameter values that were used for the model in Figure 3 are: number of managed colonies: 1,000; feral colonies: 2,000; resistance in feral and managed population at start of year 1: 5%; resistance bands [*x* = 0 to 0.8, *y* = 0.8 to 1.2, *z* = 1.2 to 2.0], probability of death [*u* = 0.8, *v* = 0.5, *w* = 0.12].

**Figure 3:**
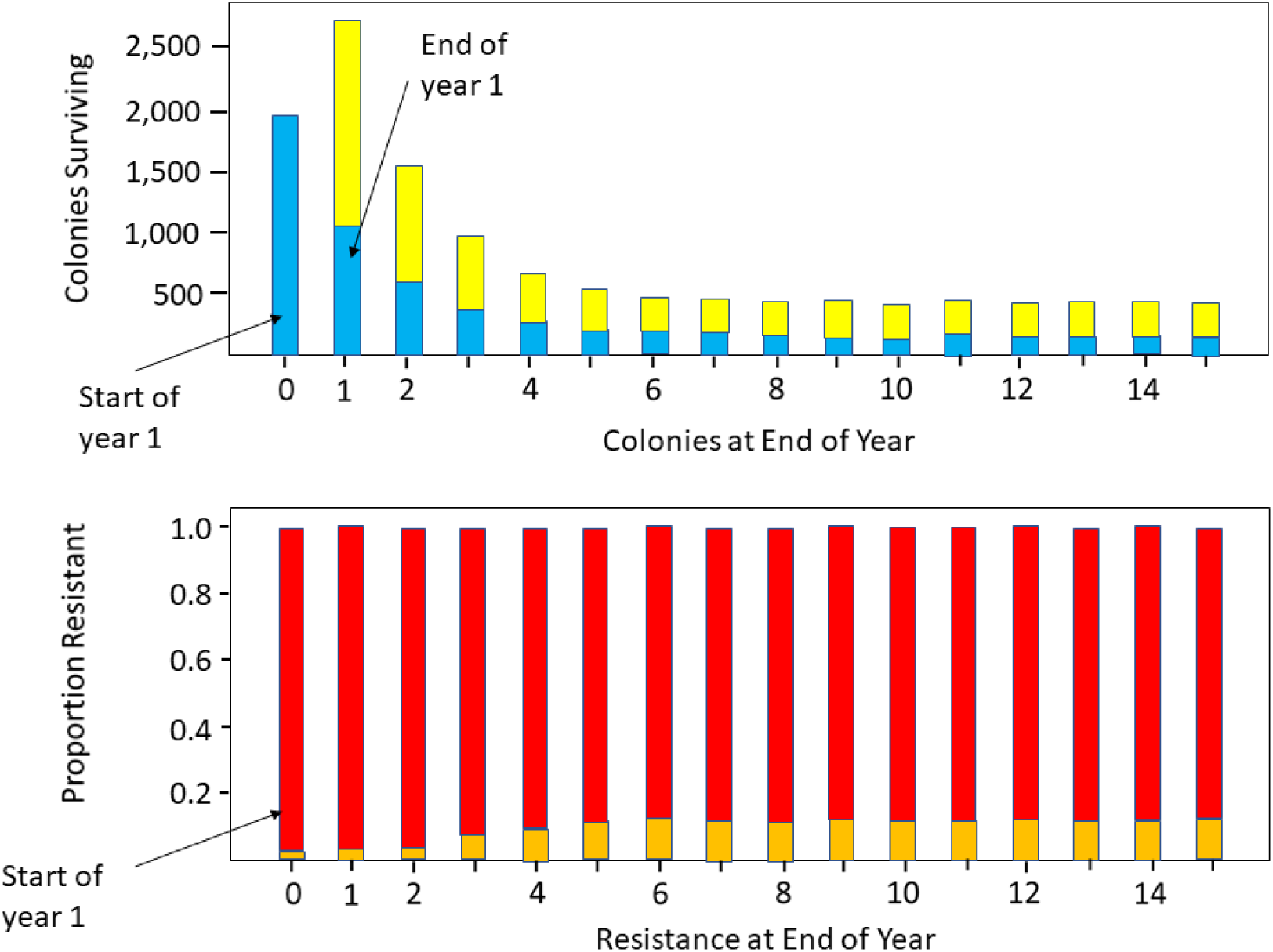
Stacked bar charts showing: Fig. (a) the number of colonies surviving 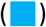, killed by *Varroa* 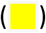 and killed by the environment 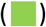 as a function of simulation year. Fig. (b) the percentage of feral colonies with high 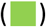, medium 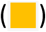 and low 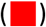 resistance as a function of simulation year. At start of year 1, 5% of feral colonies display medium resistance, 95% display low resistance. In this scenario, the probability of death by *Varroa* is high for colonies with medium levels of resistance, this causes colony death rate to be sufficiently high that the feral population shows no sign of recovery by year 15. See text for details

## Discussion

A surprising feature of our results is that introducing *Varroa*-resistant queens into the managed population had little effect on the development of resistance in the feral population. The reason for this is that the effect of selection on the feral population is fast (i.e., the feral population typically collapses within 1-3 years), while the spread of resistance from the managed population to the feral population is relatively slow. The only exception is if the size of the managed (resistant) population is large compared to the size of the feral population. This situation is not likely to be present in most areas of Australia, but may be the case locally when large numbers of apiaries are moved into certain locations to fulfill pollination contracts. Importantly, this might also be the case in urban areas where the concentration of non-commercial beekeepers can be relatively high and habitat for feral hives limited. This implies that the uptake of resistant queens by beekeepers should be as high as possible if this strategy is to have an effect on the feral population. This is also particularly important as it is likely that *Varroa* would enter Australia via one of its port-cities, inadvertently imported by shipping activities. Indeed, so far, all of the infestations detected by Australian quarantine services have been in or around shipping ports (Barry, 2010; Clifford et al., 2011; Heersink et al., 2016). Hence there is scope, as part of their biosecurity activities, for state and/or federal departments of agriculture to seed a high proportion of managed colonies in these areas with *Varroa*-resistant queens to form a ‘barrier’ to limit the spread of the mite away from its point of entry. For this strategy to work, it needs to be done well before *Varroa* enters Australia, in high-risk areas around ports, where the risk of an incursion is high. If *Varroa* were to break quarantine after most managed colonies around a port were seeded with VSH, the result could be catastrophic since the mite could spread undetected through the non-resistant feral population.

Our model shows that the main determinant of growth in resistance in the feral population is the presence of resistance within the feral population at the start of the simulation. The probability of colony death that appears to have the greatest effect on the development of population-level *Varroa*-resistance is for colonies where an intermediate number of workers possess *Varroa*-resistance.

Although we assumed a single cause of *Varroa*-resistance, it is likely that feral colonies may possess resistance conferred by unrelated traits. Studies have shown that survivor colonies in Europe possess resistance by low levels of several traits, not by a single trait (Büchler et al., 2010; Locke, 2016; Locke and Fries, 2011; Rinderer et al., 2010; Seeley, 2006; Seeley et al., 2015). However, since it is not known how effectively these different traits are co-inherited it is not unreasonable, in a first iteration of model development, to assume that they can be considered as a single trait, or several alleles located at close proximity on the same chromosome.

Within a given range of resistance bands (*x, y, z*) and probability of death (*u, v, w*), our model produces results that mirror field observations for both survivor and collapsed populations if input variables for the model are chosen appropriately. Where possible we have used realistic estimates when these were available, although many assumptions used in the model still need field validation. In fact, one of the main benefits of developing the model is that it is now much clearer which are the key parameters requiring the focus of active research (Garner and Hamilton, 2011). These include the relationship between the percentage of workers in a colony that possess resistance and the yearly probability of colony death. The spatial and temporal density of feral and managed colonies is also critical and, at the time of writing, estimates for each of these parameters are not well defined in Australia.

Parameters used in the model were varied during simulations to assess the sensitivity of model outputs. Although the proportion of the managed population seeded with resistant queens had relatively little effect on *Varroa* resistance in the feral population, it is critical that the managed population be requeened regularly with resistant queens to minimize the need to use of miticides and to allow Darwinian selection to also take place in managed honeybee populations.

To date commercial beekeepers in the USA have been reluctant to rely adopt strategies to develop *Varroa*-resistance, for example by using VSH queens (Leiby, 2014). This is partly due to concerns that VSH bees may only provide partial resistance and concerns that beekeepers with many hundreds to thousands of colonies could lose their stock of bees if resistance failed. Also, miticides are effective at managing *Varroa* at relatively low cost, even if the long-term costs may be high due to tainted honey, resistance to miticides developing in the mite and harmful effects on bees (Johnson, 2015; Johnson et al., 2015; Johnson et al., 2010; Pettis et al., 2012).

Although we have developed a model of resistance to *Varroa* we propose that it can be used to model the development of resistance to pathogens in any species that have a mating behavior similar to that of *A. mellifera*.

If Australia is to manage the long-term consequences of a *Varroa* incursion, a *Varroa*-resistant queen breeding industry needs to be developed. The use of miticides, although relatively inexpensive and effective in keeping the mite under control, results in numerous poor health outcomes for all members of the colony. For Australia to gain the most from a breeding program, it needs to be started before *Varroa* becomes endemic, not afterwards.

